# Cost and benefits of gene amplification-mediated antibiotic resistance

**DOI:** 10.64898/2026.03.19.712897

**Authors:** Yuwen Fang, Johannes Kupke, Ulrich K. Steiner, Karsten Tedin, Marcus Fulde

## Abstract

Bacteria evolve antibiotic resistance through various mechanisms, including horizontal gene transfer, sequence-altering mutations, and tandem duplications/amplifications in genomic regions that increase the dosage of resistance determinants. This phenomenon, known as gene duplication-amplification (GDA), is a transient and reversible genetic mechanism that allows bacteria to rapidly adapt to antibiotic stress. Due to the unstable nature of amplification, the quantitative relationship between gene copy number, resistance levels, and the associated physiological costs remains poorly defined. Here, we constructed strains with fixed GDA copy numbers in a clinical *Enterobacter cloacae* strain. Using growth curve and time-kill assays, we show that ceftazidime (CAZ) resistance scales with GDA copy number, but the benefit under selection comes with a growth cost in the absence of antibiotic. Notably, we find that while GDA provides a survival advantage under CAZ pressure, it simultaneously increases susceptibility to rapid killing at supra-MIC concentrations. Our analysis further illustrates that survival is jointly shaped by gene dosage, antibiotic concentration, and exposure time. Together, these results clarify that GDA-mediated resistance is highly context dependent, shaped by trade-offs between resistance benefits and physiological costs.

## Introduction

The global escalation of antimicrobial resistance (AMR) represents one of the most significant threats to modern medicine, contributing to an estimated five million deaths worldwide in 2019^1,2^. Among the most problematic pathogens are members of the *Enterobacter cloacae* complex (ECC), a group of Gram-negative, facultatively anaerobic bacteria frequently implicated in nosocomial infections, including pneumonia, urinary tract infections, and life-threatening septicemia^3^. Due to the high frequency of multi-drug resistance (MDR) within the *E. cloacae* complex, including resistance to third-generation cephalosporins such as ceftazidime (CAZ), the limited therapeutic options for ECC infections remains a critical clinical hurdle^4,5^.

While stable genetic mutations and horizontal gene transfer (HGT) are well-documented drivers of AMR, recent evidence highlights the pivotal role of gene duplication-amplification (GDA) in rapid bacterial adaptation^6,7^. GDA is a transient process involving alterations in gene copy number by either RecA-dependent or RecA-independent mechanisms^8^. GDAs can often be identified in bacterial isolates showing a heteroresistant (HR) phenotype, where a small subpopulation shows resistance to an antibiotic while the majority remains susceptible^9^. Owing to the low frequency of such resistant subpopulations, GDA-driven heteroresistance is often difficult to detect by standard susceptibility testing, and can therefore lead to treatment failure^10–14^. The intrinsic instability of amplifications has been suggested to be due to elevated bacterial metabolic burdens posed by the increased gene dosage, resulting in a rapid loss from populations in the absence of antibiotic pressure^15,16^.

Recent work has shown that the clinical isolate *Enterobacter cloacae* IMT49658-1 undergoes GDA under CAZ stress, producing tandem amplifications encompassing the β-lactamase gene *bla*_DHA-1_ and resulting in heteroresistance^17^. GDA copy number, antibiotic exposure, and lag time together shape the growth dynamics and acceleration of resistant subpopulations. Since these subpopulations are continuously reshaped by RecA-mediated recombination, GDA copy numbers fluctuate rapidly both within and between cells, complicating direct quantification of how gene dosage determines resistance levels, fitness costs, and survival.

To overcome this limitation, we generated strains carrying fixed GDA copy numbers by inducing amplification and deleting *recA*, thereby preventing GDA from ongoing homologous recombination. By integrating resistance levels with growth, survival, and competitive performance, we characterized how strains with different *bla*_DHA-1_ copy number behave across different ceftazidime concentrations. Quantitative modeling further revealed that the effects of amplification are shaped by gene copy number and CAZ concentration over time. In this way, GDA is shown to restructure the balance between resistance and physiological burden under different contexts. Taken together, our work provides a quantitative analysis of the trade-offs in GDA, elucidating the evolutionary logic of GDA as a transient yet powerful adaptive strategy under antibiotic stress, that incurs fitness costs when released from antibiotic stress.

## Results

### Establishment of strains carrying fixed GDA copy numbers

Previous whole-genome sequencing (WGS) showed that the clinical ECC isolate IMT49658-1 (WT) harbors four functional β-lactamase genes: a chromosomally-encoded AmpC β-lactamase *bla*_ACT-16_, plasmid-borne AmpC β-lactamase *bla*_DHA-1_, and class A β-lactamases *bla*_CTX-M-3_ and *bla*_TEM-1_ (Fig. 1A)^17^. Our previous study indicated that the ceftazidime heteroresistant phenotype was due to GDA of the *bla*_DHA-1_ gene^17^. To generate a genetic background in which *bla*_DHA-1_ is the only remaining β-lactamase, we sequentially deleted the *bla*_ACT-16_, *bla*_CTX-M-3_, and *bla*_TEM-1_ genes, generating a triple-deletion mutant (WT-Δ3). Although genomic analysis suggested these four genes were the primary determinants for β-Lactam resistance in this strain, it remained possible that the strain might harbor additional mutations contributing to β-Lactam resistance. To verify the absence of potential interfering mutations, we further constructed a deletion mutant (WT-Δ4), in which the *bla*_DHA-1_ gene was also deleted from the WT-Δ3 background. As shown in the susceptibility test, the WT-Δ3 strain exhibited a reduced but measurable level of resistance compared to the WT, consistent with the presence of *bla*_DHA-1_ and the loss of other β-lactamases. In contrast, WT-Δ4 displayed a complete loss of detectable resistance (Fig. 1B). This result established *bla*_DHA-1_ as the only active determinant of CAZ resistance in the WT-Δ3 background, providing a clean genetic background for examining GDA dynamics.

**Fig. 1:**
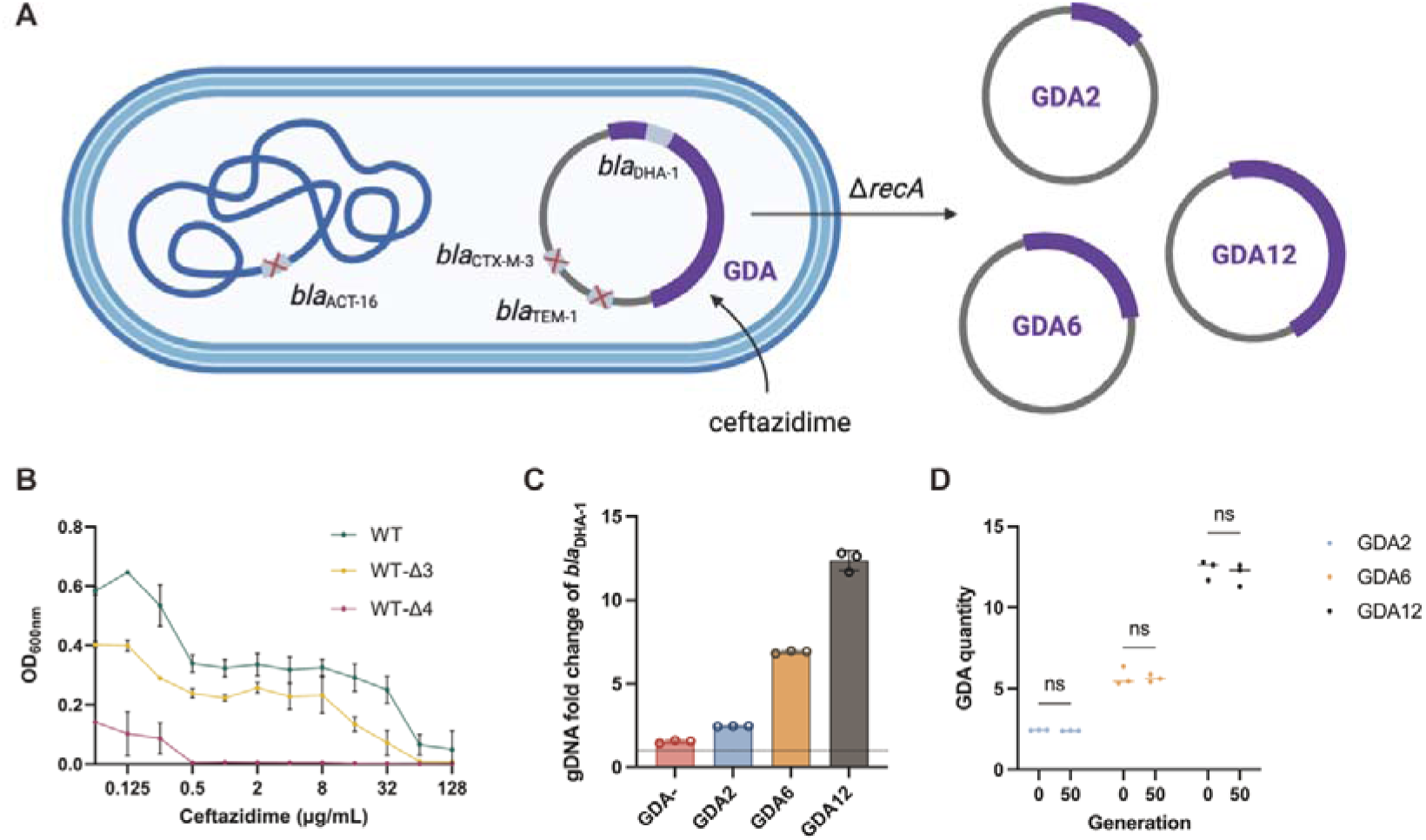
Construction and validation of strains carrying stable GDA number. **A,** Schematic diagram of the experimental strategy for generating strains with a fixed GDA copy number. The *bla*_ACT-16_, *bla*_CTX-M-3_, and *bla*_TEM-1_ genes were first deleted from the wildtype strain (WT-Δ3). Gene-duplication and amplification of the *bla*_DHA-1_ region was induced in the WT-Δ3 strain by ceftazidime (CAZ), and the *recA* gene was deleted in induced cells to prevent further homologous recombination and fix the resulting GDAs. Subsequent screening of isolated clones yielded strains harboring different numbers of GDA events. **B,** Broth microdilution assay confirming *bla*_DHA-1_ as the sole remaining determinant of CAZ resistance in WT-Δ3. The CAZ resistance was compared among the wildtype (WT) strain, WT-Δ3 (Δ*bla*_ACT-16_ Δ*bla*_CTX-M-3_ Δ*bla*_TEM-1_), and WT-Δ4 (Δ*bla*_DHA-1_ Δ*bla*_ACT-16_ Δ*bla*_CTX-M-3_ Δ*bla*_TEM-1_). **C,** qPCR quantification of *bla*_DHA-1_ copy number in four GDA-fixed strains. Copy numbers were normalized to the single-copy chromosomal reference *trp* and calibrated to the *bla*_DHA-1_ levels of the non-amplified control, WT-Δ3, set to 1. **D,** Stability of GDA copy number after ∼50 generations of serial passaging in antibiotic-free medium. *bla*_DHA-1_ copy numbers were quantified by qPCR as in panel C. n.s., not significant, determined by two-way ANOVA with Sidak’s multiple-comparisons test. Data were presented as means ± S.D from three independent biological replicates for panel B, C, D.

A major challenge in quantifying GDA physiology is that RecA-dependent recombination makes tandem amplifications highly dynamic, causing rapid fluctuations in copy number that confound direct gene-dosage measurements^18^. To capture defined amplifications, we generated strains with fixed numbers of GDA. The WT-Δ3 strain was first exposed to CAZ to enrich the subpopulation carrying the spontaneously amplified GDA region containing the *bla*_DHA-1_ gene, followed by deletion of *recA* (Fig. 1A). From the resulting *recA*-deficient strains, a total of nine independent clones were isolated. Quantitative PCR (qPCR) revealed substantial variation in *bla*_DHA-1_ copy number among these clones (Supplementary Fig. S1). To enable a comprehensive analysis of gene dosage effects, three representative strains were selected: a low-copy variant (GDA2, ∼2 copies), a medium-copy variant (GDA6, ∼6 copies), and a high-copy variant (GDA12, ∼12 copies) (Fig. 1C). In parallel, a non-amplified control strain (GDA−) was generated from WT-Δ3 without prior antibiotic exposure.

To verify the stability of these mutants, we performed experiments in which all four variants (GDA−, GDA2, GDA6, and GDA12) were serially passaged in antibiotic-free medium for ∼50 generations. As shown in Fig. 1D, no significant change in *bla*_DHA-1_ copy number was observed at the endpoint of the experiments. These results confirmed that the *recA*-deficient background effectively locked the amplification states, allowing direct comparison of strains differing only in GDA copy number.

### GDA copy number drives ceftazidime resistance

Using the established GDA-fixed strains, we first evaluated the impact of *bla*_DHA-1_ gene dosage on resistance to CAZ. Broth microdilution assays showed that resistance scaled with gene copy number. As seen in Fig. 2A, growth of the GDA− variant showed no detectable growth at 2 µg/mL, whereas the GDA2 and GDA6 strains were inhibited at 16 µg/mL and 64 µg/mL, respectively. Notably, GDA12 maintained visible turbidity up to 64-128 µg/mL.

**Fig. 2:**
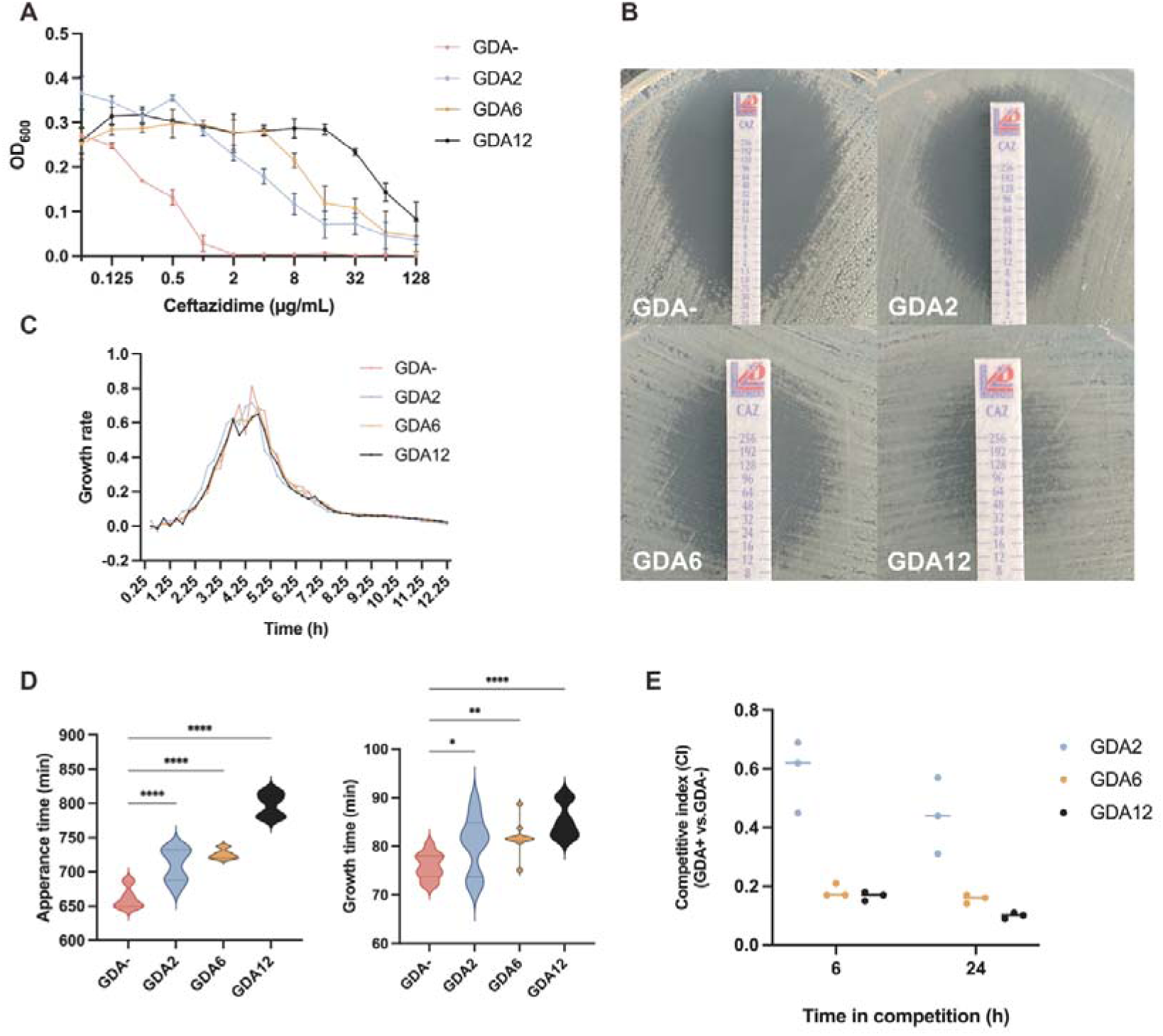
GDA copy number drives resistance at the cost of physiological fitness. **A,** Broth microdilution determinations of the resistance levels of GDA−, GDA2, GDA6, and GDA12 to ceftazidime (CAZ). **B,** E-tests with CAZ strips defining minimum inhibitory concentrations (MICs) of GDA−, GDA2, GDA6, and GDA12. The MIC values were read at the intersection of the inhibition zone and the strip scale. The MICs are 0.38 µg/mL (GDA−), 4 µg/mL (GDA2), 12 µg/mL (GDA6) and 24 µg/mL (GDA12). The absence of colonies within inhibition zones indicates a homogeneous resistance phenotype without subpopulations. **C,** Growth rates of GDA−, GDA2, GDA6, and GDA12 in antibiotic-free medium (n = 3). Growth rates were calculated as the slope of ln(OD_600_) versus time after 3-point moving-average smoothing. **D,** Appearance time and growth time detection of single colony in CAZ-free plates using ScanLag (n = 9). The dotted lines inside each violin define the interquartile range (IQR). **E,** Competition assay for GDA2, GDA6, and GDA12 against the GDA− strain (n = 3). Competitive indexes (CI) were determined at 6 h and 24 h of competition in the absence of antibiotics. CI values less than 1 indicate reduced competitiveness relative to the GDA− strain. Data in C, E were presented as mean values ± S.D. P-values in D were determined by one-way ANOVA followed by Dunnett’s multiple-comparisons test.

Examination of the optical density (OD) further revealed quantitative growth differences. The GDA− strain exhibited a sharp collapse over a narrow concentration range. In contrast, the amplified variants displayed a progressively gradual decline, with GDA12 maintaining measurable biomass even at 128 µg/mL.

E-test measurements confirmed the gene-dose effect. The WT exhibited a minimum inhibitory concentration (MIC) of 2 µg/mL, whereas the WT-Δ3 showed a reduced MIC of 0.75 µg/mL (Supplementary Fig. S2). The lowest MIC was found in GDA− at 0.38 µg/mL. Introduction of low-level amplification increased the MIC to 4 µg/mL in GDA2, with further scaled increases observed in GDA6 and GDA12, reaching 12 µg/mL and 24 µg/mL, respectively (Fig. 2B). Interpreting these values with standard CLSI breakpoints revealed the significance of amplification: while the MIC of GDA− falls within the susceptible range (≤4 µg/mL), the amplification to ∼12 copies shifts the phenotype to fully resistant (≥16 µg/mL). This represents a >60-fold MIC increase driven solely by the amplification of the *bla*_DHA-1_. Together, these results revealed a strong positive correlation between the GDA quantity and ceftazidime resistance within the tested range.

### Amplification impairs growth in non-selective conditions

Resistance acquisition often comes at a fitness cost. To quantify the physiological burden of maintaining amplified *bla*_DHA-1_, we measured growth kinetics in antibiotic-free medium. Growth rate analysis revealed a clear inverse relationship between GDA copy number and growth performance (Fig. 2C). The GDA− strain rapidly reached its maximum growth rate, whereas the high-copy variants exhibited lower peak growth rates. In addition, the maximum growth rate progressively declined with increasing GDA copy number. Furthermore, GDA12 strain accumulates the least biomass (OD) after 24 h (Supplementary Fig. S3). These observations suggest that the amplified *bla*_DHA-1_ gene, and potentially other genes within the GDA region, confer growth costs.

To further assess this growth defect, ScanLag^19^ was performed to track both the appearance time and growth time of individual colonies (Supplementary Fig. S4). The results confirmed that GDA significantly delays both the colony appearance and subsequent colony growth. Specifically, colony appearance time, defined as the time for a colony to reach a detectable threshold of 10 pixels, was delayed by ∼64 min in GDA6 and ∼137 min in GDA12 compared with GDA− (Fig. 2D), indicating a prolonged lag phase and slowed early growth. Crucially, this growth defect persisted after colony emergence. The mean growth time, measured as the duration to expand the colony area from 15 to 45 pixels, was significantly prolonged in the amplified variants, with GDA12 exhibiting a ∼9.4 min increase compared to GDA−, representing a ∼12.4% increase. Together, these results demonstrate that amplification not only prolongs the lag phase but also imposes a continuous physiological burden during the exponential phase.

Competition assays in antibiotic-free medium further demonstrated the fitness disadvantage. When co-cultured with the GDA− strain in antibiotic-free medium, amplified strains were rapidly outcompeted (Fig. 2E). Within 6 hours, competitive indices (CI) dropped to ∼0.17 for GDA12 and ∼0.18 for GDA6, indicating that the fitness impairment manifests almost immediately at the start of co-culture. By 24 hours, GDA12 declined further to a CI of ∼0.10. Even the low-copy GDA2 strain showed a gradual disadvantage (CI 0.59 at 6 h, 0.44 at 24 h). These data support a evolutionary trade-off: while amplification increases resistance, it imposes an immediate and substantial fitness cost under non-selective conditions^13^.

### Trade-offs between growth and survival across antibiotic gradients

To systematically assess the trade-offs shaped by gene amplification, we analyzed bacterial performance across different CAZ concentrations. The growth curves and the analysis of growth dynamics (area under the curve, AUC) confirmed a striking inversion of fitness cost as antibiotic pressure increased (Fig. 3A; Supplementary Fig. S5). At low CAZ concentrations, the physiological cost of amplification dominated the phenotype. GDA2 and GDA6 exhibited superior growth kinetics compared to GDA12, characterized by higher growth rates and/or greater biomass accumulation. For instance, at 2 µg/mL CAZ, the GDA12 showed growth deficiency, with an AUC reduction of ∼6.4% relative to GDA2. This mirrors our previous findings that under low-stress environments, the physiological burden of maintaining a high copy number of GDA outweighs the potential benefits, putting high-GDA strains at an evolutionary disadvantage. As ceftazidime concentrations increased, this relationship progressively inverted. The fitness crossover occurred between 2 and 4 µg/mL, beyond which the protective effect of *bla*_DHA-1_ amplification outweighed its cost. At intermediate and high concentrations, growth of the low-GDA strains collapsed, whereas the high-copy variants continued to grow. For example, at 32 µg/mL CAZ, the GDA− strain was almost completely suppressed (AUC ∼2.4, indistinguishable from the baseline), while GDA12 retained ∼80% of its maximum growth potential (AUC ∼13.8). This represents a strong selective advantage, indicating that high gene dosage is indispensable in a high-stress environment.

**Fig. 3:**
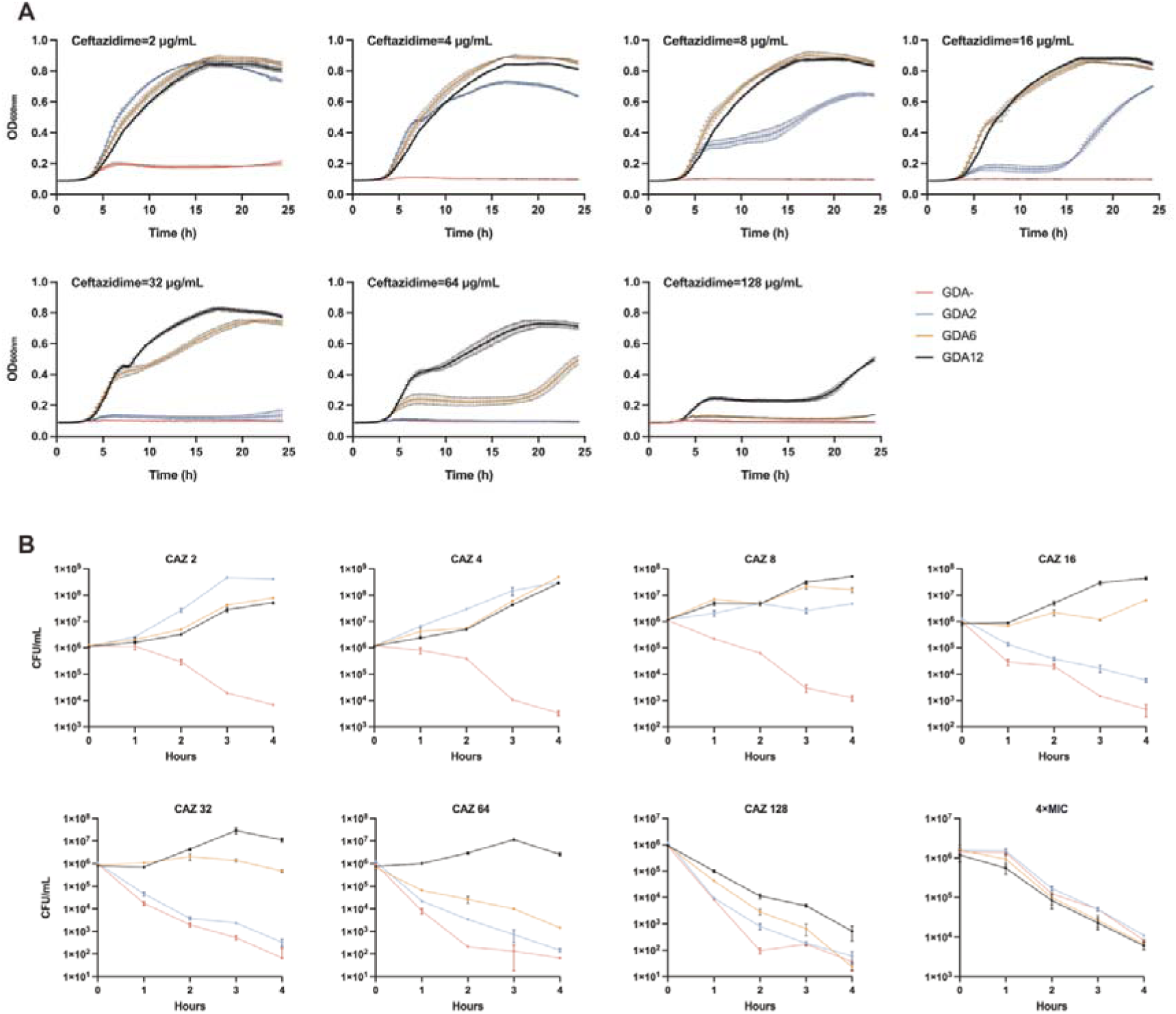
GDA copy number shapes the trade-off between growth advantage and survival under antibiotic stress. **A,** Growth curves of the GDA−, GDA2, GDA6, and GDA12 strains cultured under different CAZ concentrations (2, 4, 8, 16, 32, 64, 128, and 256 μg/mL). **B,** Time kill assay of the GDA−, GDA2, GDA6, and GDA12 strains over 4□h at different ceftazidime concentrations (2–256 μg/mL), and at 4□×□the MIC determined for each strain. Colony-forming unit counts (CFU/mL) at each time point quantify bacterial survival. Data are shown as means ± S.D from three independent biological replicates for all panels.

Since optical density captures signal from both live and dead cells, we next performed time-kill assays across the same concentration range to evaluate how GDA amplification impacts bacterial survival (Fig. 3B). At low CAZ concentrations, all strains exhibited population expansion over time, although GDA12 expanded more slowly than GDA2, consistent with the growth kinetics results. With increasing CAZ concentrations, the effect on GDA− and GDA2 rapidly transitioned to killing. In these backgrounds, CFU counts declined sharply within the first 2-3 hours, frequently exceeding a 3-log_10_ reduction, indicating bactericidal activity. In contrast, GDA12 consistently maintained high viable counts across intermediate and high CAZ concentrations, with CFU levels remaining stable or increasing over time, indicating effective resistance to antibiotic killing. GDA6 showed an intermediate phenotype, with partial killing at higher concentrations but substantially delayed and reduced CFU loss compared to GDA− and GDA2. Notably, at concentrations where the growth kinetics suggested almost complete inhibition of low GDA strains, time-kill assays revealed rapid loss of survival, whereas high-copy variants retained substantial populations. These results demonstrated that increased *bla*_DHA-1_ dosage not only permits residual growth under high antibiotic stress but also alters killing dynamics across concentrations, allowing the bacterial population to avoid complete eradication.

While GDA12 effectively maintained viability over a wide range of antibiotic pressures, it remained unclear whether this high resistance level also implies increased tolerance, defined as enhanced survival during transient exposure to lethal drug levels. Therefore, we performed time-kill assays at standardized supra-MIC conditions (4× the MIC for each strain; MIC differed substantially among the strains). Unexpectedly, despite their high resistance levels, strains carrying high GDA copy numbers were killed more rapidly than low-copy strains under these high MIC conditions. While the GDA− strain declined slowly at its 4× MIC (1.5 µg/mL), the GDA12 strain showed a rapid loss of viability at 96 µg/mL. Because tolerance would manifest as slower killing at a fixed supra-MIC multiple (i.e., longer survival despite lethal exposure), these data indicate that increased GDA copy number primarily raises the inhibitory threshold (MIC) rather than increasing tolerance to lethal exposure.

Together, our data demonstrate that GDA amplification creates a dose-dependent trade-off: high GDA copy numbers permit growth and survival across a broad range of antibiotic concentrations but do so at the cost of resilience toward acute stress: once the threshold is exceeded, high-copy variants undergo accelerated killing.

### Quantitative modeling reveals trade-offs between resistance and fitness associated with GDA copy number

To predict responses, we parameterized models with the observed empirical data. We first examined the relationship between *bla*_DHA-1_ copy number and antibiotic resistance (Supplementary Table S1). Model selection based on Akaike Information Criterion (AIC) indicated that, within the tested range (1-12 copies), gene dosage acts as a direct and scalable determinant of MIC. We next quantified the fitness cost of GDA using ScanLag measurements (Supplementary Table S2). Linear regression identified GDA copy number to predict colony appearance time, demonstrating that the cost imposed by amplification directly scales with copy number. Importantly, this cost is independent of antibiotic exposure, indicating that amplification imposes a constraint on cellular capacity to grow, rather than a conditional defect expressed only under stress.

To capture the trade-offs observed across antibiotic gradients, we applied generalized additive models (GAMs) to the growth-curve data (Supplementary Table S3). Comparison of additive and interactive model parameters revealed overwhelming support for a model incorporating a GDA*concentration interaction (ΔAIC = 2756.9). This indicates that gene dosage and antibiotic pressure jointly reshape growth dynamics rather than acting independently in an additive only way. Consistent with our experimental observations, the model accurately predicted the fitness crossover, where high-GDA variants showed a growth disadvantage at low CAZ concentrations but gaining a pronounced advantage above ∼4 µg/mL.

As these interactions describe growth behavior and do not directly capture survival outcomes, we extended our analysis to the results of time-kill assays (Supplementary Table 4). A three-way interaction model incorporating time, GDA copy number, and CAZ concentration provided the best fit, indicating that killing kinetics cannot be decomposed into simple additive effects. Notably, the model captured the observed trade-off whereby high-GDA strains exhibited enhanced survival at moderate concentrations but underwent accelerated killing once supra-MIC levels are exceeded.

Overall, the fitted models closely recapitulated the experimental trends (Supplementary Fig. S6-9). To verify the model predictions, we compared simulated growth curves as predicted by the model and empirical growth curves from an independently constructed strain carrying an intermediate GDA copy number (∼5 copies; GDA5) (Fig. 4). Across the CAZ gradient, the predicted curves closely recapitulated the overall shapes of the experimental growth curves, capturing key features such as progressive suppression of growth as antibiotic concentration increased. Deviations were the most evident near the inhibition concentration, where residual growth becomes small and more variable. Together, these results indicate that these models provide a quantitative, generalizable description of how GDA copy number, CAZ concentration and exposure time jointly determine resistance, growth and killing outcomes.

**Fig. 4:**
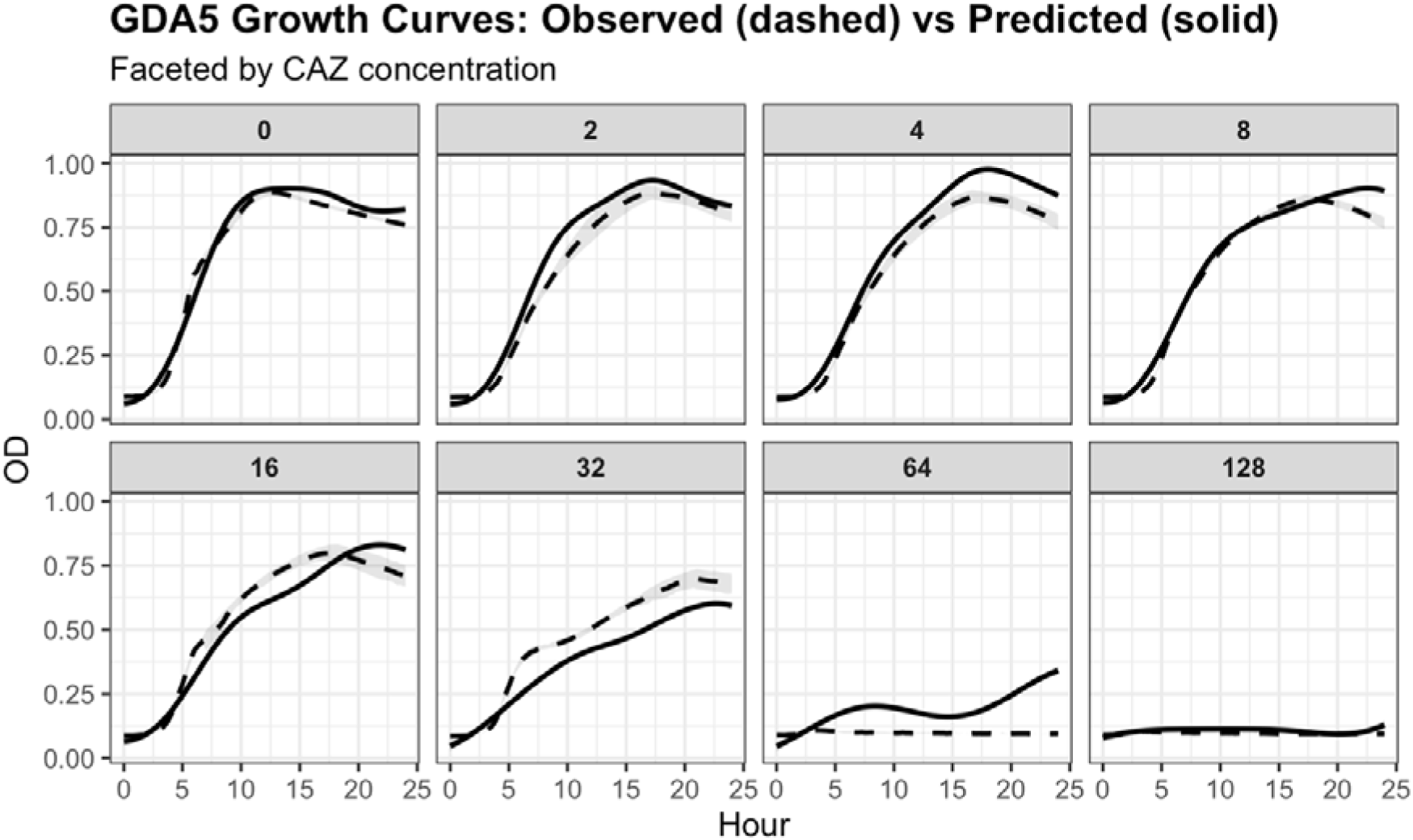
Model-based prediction of growth dynamics for a ∼5-copy strain (GDA5). Model-based predictions for an independently constructed strain (GDA5, ∼5 copies) across ceftazidime (CAZ) concentrations. Solid curves show the growth curves predicted by the AIC-selected growth-curve model (GCGam3c) trained on the main dataset (GDA−/GDA2/GDA6/GDA12), using measured copy number and CAZ concentration as inputs without refitting. Dashed curves show experimentally observed mean OD_600_ for GDA5 (n = 3). Shaded ribbons indicate 95% confidence intervals for model predictions (solid; derived from model standard errors) and for observed means (dashed; calculated from replicate variability, mean ± 1.96×SE).

## Discussion

Heteroresistance can evade standard susceptibility testing and has been linked to therapeutic failure, making it a clinically important but often underappreciated resistance phenotype^9^. Gene duplication-amplification has emerged as a primary driver of heteroresistance, particularly in Gram-negative bacteria, providing a rapid and plastic response to antibiotic pressure^20^. In this study, we quantified the trade-off between the resistance benefits and fitness costs conferred by amplification of the β-lactamase gene *bla*_DHA-1_ in an ECC isolate. By constructing strains with fixed GDA numbers, we directly measured, for the first time, how amplification copy number affects antibiotic resistance, growth, and survival in various contexts. Our findings reveal a double-edged trade-off for β-lactamase amplification: increasing gene dosage proportionally increases antibiotic resistance, but at the cost of significant physiological burden, and the net effect on fitness is highly dependent on the antibiotic environment.

A clear dose-dependent relationship between gene copy number and resistance level was observed, consistent with earlier reports that heterogeneous resistance in bacterial populations can often be explained by variable gene dosage^21^. Recent ultra-deep long-read sequencing work by Jonsson et al. further supports this concept by showing that the heteroresistance phenotype of an *Escherichia coli* population is closely linked to the distribution of tandem amplification copy numbers of a β-lactamase gene, and that this distribution shifts dynamically under antibiotic selection and upon its removal^18^. Notably, within the range we tested, the resistance gain per additional copy showed no sign of saturation. However, it is conceivable that beyond a certain point, diminishing returns would set in. For instance, if enzymatic degradation of antibiotic becomes limited by substrate availability, or if the periplasmic space becomes saturated with enzyme^22^. Indeed, other studies have noted that extremely high copy numbers sometimes confer little further MIC increase once other biological limits are reached^23^. In our case, the >60-fold increase in MIC compared to the non-amplified control achieved by 12 copies of *bla*_DHA-1_ was not enough to reach such a plateau, suggesting the linear benefit extended at least to this level of amplification.

Balanced against the resistance benefit is the fitness cost of carrying and expressing extra genes. We found that higher *bla*_DHA-1_ copy numbers severely impaired growth in antibiotic-free conditions, as evidenced by slower growth rates, prolonged lag phases, and reduced competitive fitness. This provides direct support to the long-suspected notion that heteroresistance comes at a cost^16^. Our quantitative data are in line with previous measurements of amplification costs in other bacteria. For example, in a study of heteroresistant *E. coli* and *Salmonella enterica*, different amplified regions were estimated to incur a 0.02%-0.12% fitness reduction per extra kilobase of DNA^13^. The burden likely arises from multiple factors: the energetic cost of replicating and segregating the extra DNA, the resource cost of transcribing and translating the amplified genes, as well as any disruptive effects of the overexpressed proteins on cellular physiology^24^. In our strains, overproduction of *bla*_DHA-1_ could deplete ribosomal capacity and divert amino acids and energy towards enzyme production and export, thereby delaying cell cycle progression and reducing growth rate^25–27^. These costs may be even more apparent if the co-amplified gene does not have a beneficial function under normal conditions. As a result, in the absence of antibiotics, high GDA cells show an impaired state compared to low GDA cells.

One of our most striking observations was how the selection advantages reverse as the antibiotic concentration changes. In antibiotic-free medium, high-GDA clones are almost certain to be outcompeted by the wild-type (or low-GDA) population, explaining why amplified subpopulations typically decrease once antibiotic stress is relieved^28,29^. However, under increasing antibiotic stress, there is a threshold concentration at which the amplified subpopulation starts to outperform the others. This threshold can be understood as the point where the resistance advantage conferred by amplification outweighs its fitness cost — essentially the minimal selective concentration (MSC) for the amplified variant^30,31^. Below the MSC, amplification is a liability; above it, amplification becomes an advantage, and the high-copy subpopulation will be selectively enriched. Our data indicate the MSC for the ∼12-copy strain was ∼4 µg/mL CAZ, which is about 4-fold below its MIC, and just above the MIC of the susceptible strain. This finding aligns with theoretical expectations that higher-cost resistance mechanisms require higher antibiotic levels to be favored^32^. High-cost amplifications like our GDA12 strain demand a relatively high antibiotic concentration to yield a positive selection coefficient, whereas a lower-cost amplification can be favored at a lower concentration. This dynamic explains why heteroresistance is often transient: when antibiotic concentrations fluctuate (such as during dosing cycles or varying tissue penetration), the advantage of amplification may come and go, causing continuous fluctuations in the resistant subpopulation’s frequency.

Our work also reveals a paradoxical vulnerability of the high-copy strains: reduced tolerance to extreme antibiotic stress. While GDA12 survived and grew at CAZ levels that killed GDA−, once the concentration greatly exceeded the MIC of GDA12, cells were killed more rapidly than those with lower GDA levels. In essence, amplification raised the level of resistance but did not protect, and apparently sensitized, the cells against overwhelming drug exposure. This highlights the distinction between resistance, defined as the ability to grow at elevated antibiotic concentrations, and tolerance, defined as the ability to survive lethal exposure for limited periods of time^33^. Our GDA12 strain had high resistance but low tolerance in the sense that it was rapidly killed at 4× MIC. A plausible explanation is that, at supra-MIC concentrations, β-lactams trigger toxic dysregulation of cell wall biosynthesis while also promoting metabolic remodeling and oxidative stress^34–37^. In high-GDA strains, the pre-existing burden of gene amplification may further limit stress-buffering capacity, making them disproportionately susceptible to rapid killing under saturating ceftazidime exposure.

An important aspect of our study was the introduction of a *recA* mutation to stabilize gene amplifications. RecA-mediated homologous recombination is known to drive both the formation and collapse of tandem duplications^38,39^. The strategy of removing *recA* allowed us to treat gene copy number as a fixed variable, revealing its contributions unmasked by ongoing recombination. The approach is similar in spirit to recent research which used recombination-deficient mutants to study amplification in *E. coli*^40^. Heteroresistant isolates often display colonies growing within the zone of inhibition in the E-test, reflecting small resistant subpopulations within a susceptible culture. In our *recA*-deleted GDA strains, no such heteroresistant colonies were seen, confirming that their phenotype was homogeneous and stable. While secondary, *recA*-independent pathways may exist^23,41^, their contribution to genomic rearrangement in our study appears negligible compared to the stability afforded by *recA* inactivation.

These findings have several implications for the treatment of ECC infections in clinical settings. By quantifying the cost of GDA, our study explains why amplification, despite its frequent emergence, is typically replaced by mutations in the long term^42^. Once a stable beneficial mutation emerges, maintaining additional copies are often no longer necessary and may become a burden^43^. Notably, as antibiotic concentrations increase, the required chromosomal alterations typically become more complex, inherently carrying significant physiological costs that may eventually parallel the burden of gene amplification. Moreover, the vulnerability of high-GDA strains at supra-antibiotic concentrations suggests potential therapeutic opportunities, provided these required doses remain within clinically safe limits for the patient. If these high-copy bacteria exist in a metabolically stressed state, they may be particularly susceptible to agents targeting the membrane or related pathways. Combination therapies may therefore trap bacteria in an evolutionary dilemma^44^.

This study has some limitations. By using a *recA* deletion mutation to stabilize amplifications, we created an artificial scenario to measure effects in isolation. In real infections, amplifications will not be static but dynamically changing. However, the trends we observed should still apply qualitatively to those transient amplifications. Furthermore, our work focused on a single resistance gene (*bla*_DHA-1_) and antibiotic (ceftazidime). Different resistance mechanisms or different antibiotics may have distinct cost-benefit landscapes. For instance, an amplified efflux pump might have a different metabolic cost or might confer cross-resistance to multiple drugs, altering the dynamics. Additionally, our experiments were performed in a rich growth medium. In nutrient-limited or host environments, the metabolic cost of GDA may be higher, or alternatively, be masked by growth limitations. These factors warrant further investigation. Finally, while our study revealed the trade-offs in heteroresistance, the molecular basis of the costs remains to be elucidated. Exploring factors such as cellular systems which become limiting, and which stress responses are triggered by amplification may reveal new targets against heteroresistant pathogens. In summary, amplification-mediated resistance is highly context dependent and is jointly determined by gene dosage, antibiotic concentration and exposure time, with benefits that can reverse under supra-MIC exposure.

## Materials and methods

### Strains, Media, and Antibiotics

The bacterial strain used as the parental background in this study (IMT49658-1 Δ*catA2*) has previously been described^17^. The strain was derived from a clinical *Enterobacter cloacae* complex (ECC) isolate IMT49658-1 (*Enterobacter hormaechei*), originally isolated from an equine wound infection at the Freie Universität Berlin (NCBI BioProject: PRJNA1020684).

Bacteria were cultured using cation-adjusted Mueller-Hinton II (MH II) broth or agar (Becton Dickinson) for antibiotic resistance testing, and in Lysogeny broth (LB) (Lennox formulation; Carl Roth) for all other experiments. Unless noted otherwise, antibiotics were used at the following concentrations for selection during mutagenesis: hygromycin (Carl Roth; 75 µg/mL), phleomycin (InvivoGen; 15 µg/mL), and chloramphenicol (Sigma-Aldrich; 30 µg/mL). Neomycin (Carl Roth; 50 µg/mL) was used for plating in the competition assays. Ceftazidime (CAZ) (Sigma-Aldrich) was used when comparing the phenotypes of the GDA-fixed strains at the concentrations indicated in the text.

### Targeted Gene Deletion Mutagenesis

Targeted gene deletions (Δ*bla*_ACT-16_, Δ*bla*_CTX-M-3_, Δ*bla*_TEM-1_, Δ*bla*_DHA-1,_ and Δ*recA*) were performed using the λ-Red recombinase system essentially as described^45^, with modifications. To generate deletion mutants, electrocompetent cells were prepared from IMT49658-1 Δ*catA2* harboring pSIM18^46^, encoding the hygromycin resistance gene and a heat-inducible λ-Red recombinase. Cultures were grown at 29°C to an optical density at 600 nm (OD_600_) of approximately 0.3. Cultures were induced at 42°C in a water bath for 15 min to induce the recombinase, followed by immediate cooling on ice. Cells were washed three times by centrifugation and resuspension in ice-cold 10% glycerol.

Mutagenic DNA fragments were amplified via PCR (Biometra T3000 PCR Thermocycler) using the template plasmid pKD4ble, a derivative of pKD4 carrying a bleomycin/phleomycin resistance cassette flanked by FRT sites. PCR primers contained ∼50 bp homology extensions flanking regions of the target genes. Purified PCR products (∼330 ng) were electroporated into the induced cells using a MicroPulser Electroporator (Bio-Rad) Ec2 program (V = 2.5 kV). After recovery in SOC medium at 37°C, 200 rpm for 1 h, transformants were selected on LB agar containing phleomycin. Putative gene deletion clones were verified by PCR using gene-specific external and bleomycin-specific internal primer pairs (see Supplementary Table S5), and sequencing confirmation of the deletions.

Competent cells were prepared from the different gene deletion mutant strains and the antibiotic resistance cassette was excised by introducing the FLP recombinase-expressing plasmid pCP20. Loss of the cassette was verified by phleomycin sensitivity tests and PCR. FLP plasmid curing was then achieved by growth at 37°C, followed by screening for loss of chloramphenicol resistance at 30°C.

### Quantification of GDA Copy Number via qPCR

Copy number variations of the resistance gene *bla*_DHA-1_ were determined using quantitative PCR (qPCR). Genomic DNA was extracted using QIAamp DNA Mini Kit (Qiagen), and the concentration was assessed with NanoDrop 1000 Spectrophotometer (Thermo Fisher Scientific).

qPCR was performed using 6.7 ng DNA per reaction with Power SYBR Green Master Mix (Thermo Fisher Scientific) in 96-well plates (Eppendorf twin.tec® PCR Plate 96, unskirted, white) and analyzed on PCR cycler StepOnePlus^TM^ (Thermo Fisher Scientific). Cycling conditions consisted of an initial denaturation at 95 °C for 10 min, followed by 40 cycles of 95 °C for 15 s, and 60 °C for 1 min. A melt curve analysis was performed immediately after cycling, with 95°C for 15 s, 60°C for 1 min, and 95°C for 15 s.

The single-copy housekeeping gene *trp* (tryptophan synthase) was used as the chromosomal reference for normalization. Relative copy number was calculated using the 2^−ΔΔCt^ method^47^. Fold-changes were determined by comparing the ΔCT values of the GDA-carrying strains to those of the susceptible, non-amplified parental strain, which was set to 1 copy. The primer sequences used in the qPCR analysis are listed in Supplementary Table S5.

### Stability Analysis of GDA Copy Number

To assess the genetic stability of the GDA in the *recA*-deficient background, serial passaging was performed in the absence of antibiotic selection. The three *recA*-deficient GDA strains (GDA2, GDA6, GDA12) and the non-amplified control (GDA−) were initially cultured overnight in antibiotic-free LB broth at 37°C. For each strain, the overnight culture was subcultured daily into fresh antibiotic-free LB broth at a dilution of 1:1000 (∼10 generations per passage). Following the final passage, genomic DNA was extracted, and the copy number of the region containing *bla*_DHA-1_ was quantified via qPCR as described above to determine if the amplification levels remained fixed or reverted to the wild-type state.

### Antimicrobial Susceptibility Testing

#### Broth Microdilution

Antimicrobial susceptibility was first characterized using the standard broth microdilution assay according to the CLSI 2020 guideline. Log-phase cultures (OD_600_ = 0.5) were adjusted to 2 × 10^6^ CFU/mL in MH II broth and mixed with equal volumes of two-fold serial antibiotic dilutions in 96-well flat-bottom plates (Greiner Bio-One) and incubated at 37°C. At 0 h and 20 h, OD_600_ was measured using Synergy HTX Multimode Reader (BioTek). The displayed OD values represent the OD at 20 h minus the baseline OD at 0 h.

#### E-tests

Minimum inhibitory concentrations (MICs) were determined using E-test strips (Biomérieux) containing CAZ. Bacterial suspensions were adjusted to a McFarland standard of 0.5 in NaCl, then swabbed onto MH II plates and incubated for 16-20 h at 35°C according to CLSI VET01 (performance standards for Antimicrobial Disk and Dilution Susceptibility Tests for Bacteria isolated from Animals).

### Growth Kinetics

Bacterial suspensions were adjusted to 1 × 10^6^ CFU/mL in LB broth. A volume of 100 µL was added in 96-well flat-bottom plates (Greiner Bio-One). Growth curves were established by OD_600_ determination with an interval of 15 min for 24 h at 37 °C using Synergy HTX Multimode Reader (BioTek).

### ScanLag Analysis

ScanLag experiments were performed to quantify single-colony lag time and growth time, following the methodology described by Levin-Reisman et al.^19,48^. Bacteria grown to exponential phase (OD_600_ = 0.5) were diluted to 10^3^ CFU/mL and plated on MH II agar to obtain 10-50 colonies per plate. Plates were covered with black felt and incubated at 33°C in an incubator outfitted with EPSON Perfection V370 scanners. Images were acquired every 15 min for 24 h using UnixScanningManager.

Image analysis was conducted in MATLAB R2020a using a custom script based on the NQBMatlab software, as previously described^17^. Two growth parameters were extracted: colony appearance time (lag time), defined as the time required for a colony to reach a detection threshold of 10 pixels, and the growth time, defined as the time required for colony area expansion from 15 to 45 pixels. Colonies that merged during incubation were identified and right-censored to ensure analytical accuracy. Merging events were mathematically defined as colonies exceeding 30 pixels in size that exhibited a sudden area increase of >1.5-fold within 15 min. Statistical analyses and data visualization were performed using R (R Core Team), with plots generated using the ggplot2 package.

### Competition Assays

Pairwise competition assays were performed between the GDA− and GDA+ strains (GDA2, GDA6, and GDA12) in antibiotic-free LB broth. Competing strains were distinguished using the plasmid pProbe′-gfp(LAA) (Addgene cat. no. 40171), which encodes the red fluorescent protein mScarlet (derived from pMRE135) and carries a neomycin resistance cassette. Cultures were mixed at a 1:1 ratio in 6 mL LB (2 × 10^6^ CFU/mL, 200 µL per strain) and incubated at 37°C, 200 rpm. Population ratios were determined at 0, 6, and 24 h by washing aliquots in PBS and plating on selective and non-selective agar. Competitive indices (CI) were calculated to quantify the relative fitness of GDA+ strains compared with the GDA−. The input represents the ratio of CFU counts of GDA to GDA− strains at the start of the competition (t = 0), while the output represents the ratio measured at the indicated time points after co-culture. CI was calculated using the following formula:

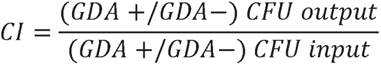

### Time-Kill Assays

Time-kill assays were conducted in 6 mL LB broth inoculated with exponential-phase cultures (OD_600_ = 0.5) to achieve a final concentration of approximately 1 × 10^7^ CFU/mL. For standardized killing assays, ceftazidime (CAZ) was added at 4× MIC of each strain, as previously determined in the E-tests. For gradient assays, a range of CAZ concentrations was applied. Bacterial survival was monitored by sampling at hourly intervals (0, 1, 2, 3, and 4 h), followed by 10-fold serial dilution and plating on MH II agar. Plates were incubated at 37 °C overnight. Viable counts (CFU/mL) were determined and plotted against time.

### Statistical modeling and model selection

Analyses were performed in R. For MIC, ScanLag, growth-curve and time-kill datasets, we fitted sets of competing models and selected the best-supported model using Akaike Information Criterion (AIC). Candidate model structures and comparison statistics are reported in Supplementary Table S1-S4. For the independent validation strain (GDA5), model-based predictions were generated from the AIC-selected GAM using the measured copy number and antibiotic concentrations as inputs, without refitting the model. For model-based predictions, 95% confidence intervals were derived from model-based standard errors (±1.96×SE). For observed growth curves, 95% confidence intervals were calculated from replicate variability (mean ± 1.96×SE). Plots were generated in R using ggplot2.

### Statistical Analysis

Experimental workflow schematics were created using BioRender. With the exception of statistical analyses described in the ScanLag experiments, the modeling and its associated data visualization were performed using R with the ggplot2 package, while all other statistical analyses and data visualization were performed using GraphPad Prism (version 10.4.2). For the serial passage experiments, differences in *bla*_DHA-1_ copy number before and after passaging for each strain were analyzed using two-way ANOVA, followed by Sidak’s multiple-comparisons test. Final statistical analysis for the ScanLag experiments was performed using one-way ANOVA, comparing each strain with the GDA− strain. For growth curves conducted under antibiotic-free conditions, growth rates were calculated after 3-point moving-average smoothing and ln transformation, presented as the slope of ln(OD_600_) versus time. Growth curves under different CAZ concentrations were analyzed by comparing the area under the curve (AUC). All data are presented as the mean ± standard deviation (SD) from three independent biological replicates, each comprising three technical replicates. A P value of < 0.05 was considered statistically significant.

## Acknowledgments

This work was supported by the DFG Priority Programme SPP2225 (FU1027/4-1) and the Collaborative Research Center CRC1449 (Project ID 431232613, Project B5), both awarded to M.F. Y.F. was supported by the China Scholarship Council (CSC) Program. U.K.S. was supported by the Heisenberg Programme of the German Research Foundation (grant 430170797). J.K. received financial support through a scholarship from the H. Wilhelm Schaumann Foundation.

## Author contributions

Conceptualisation and study design: Y. Fang, J. Kupke, M. Fulde, K. Tedin, U. K. Steiner. Performance of experiments: Y. Fang. Statistical modelling: U. K. Steiner, Y. Fang. Writing of manuscript: Y. Fang, J. Kupke, K. Tedin, U. K. Steiner, M. Fulde. Funding acquisition: M. Fulde.

## Competing interests

The authors declare no competing interests.

## Notes

### Competing Interest Statement

The authors have declared no competing interest.

