## Supplementary material for "Cost and benefits of gene amplification-mediated antibiotic resistance"

**Content list**

Supplementary Fig. S1. qPCR quantification of *bla*$_{DHA-1}$ gene copy number in different variants.
Supplementary Fig. S2. E-test results of ceftazidime susceptibility for WT and WT-3Δ strains.
Supplementary Fig. S3. Growth kinetics of GDA−, GDA2, GDA6, and GDA12 strains in antibiotic-free medium.
Supplementary Fig. S4. Representative ScanLag analysis output for GDA−, GDA2, GDA6, and GDA12 strains under antibiotic-free conditions.
Supplementary Fig. S5. Area Under the Curve (AUC) analysis of bacterial growth under increasing antibiotic concentrations.
Supplementary Fig. S6. Gene dosage shapes MIC distributions across antibiotic concentrations.
Supplementary Fig. S7. Distributions of ScanLag-derived colony growth parameters across GDA copy-number variants.
Supplementary Fig. S8. Growth-curve dynamics across CAZ gradients and GDA copy-number variants.
Supplementary Fig. S9. Time-kill kinetics across CAZ gradients and GDA copy-number variants.
Supplementary Table S1. Model selection and comparison statistics for models fitted to Minimum Inhibitory Concentration (MIC) data.
Supplementary Table S2. Model selection and comparison statistics for models fitted to colony appearance time (ScanLag) data
Supplementary Table S3. Model selection and comparison statistics for generalized additive models (GAMs) fitted to growth curve data.
Supplementary Table S4. Model selection and comparison statistics for models fitted to time-kill kinetics data.
Supplementary Table S5. Primers used in this study.

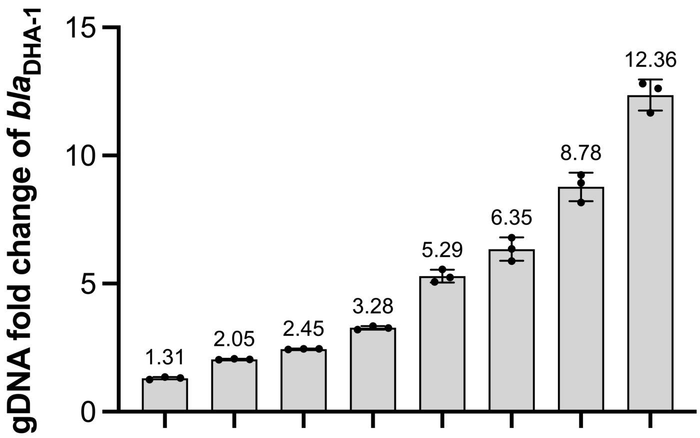


**Supplementary Fig. S1: qPCR quantification of $\textit{bla}_{\textbf{DHA-1}}$ gene copy number in different variants.** The y axis shows the $\textit{bla}_{\text{DHA-1}}$ copy number in nine independent clones obtained after ceftazidime enrichment and *recA* deletion. Values are shown as fold change relative to the non-amplified control (set to 1) and are ordered by increasing copy number. Values above the bars indicate the mean fold change. Data were presented as means ± S.D from three independent biological replicates.

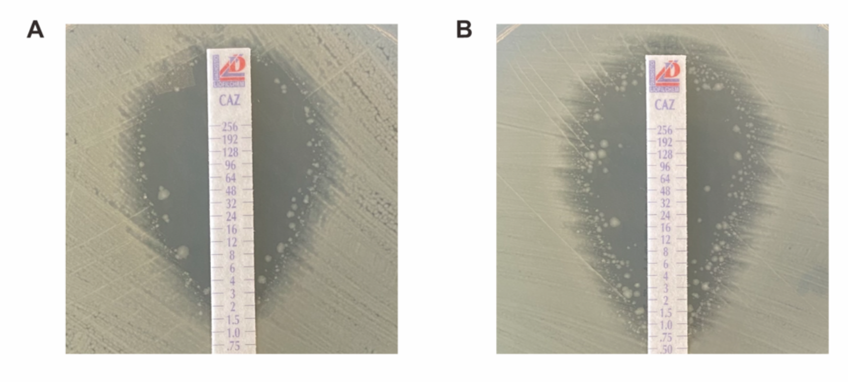


**Supplementary Fig. S2: E-test results of ceftazidime susceptibility for WT and WT-3Δ strains.** Representative images of E-test assays showing the inhibition zones for **(A)** the WT strain, and **(B)** the WT-3Δ ($\Delta \textit{bla}_{\text{ACT-16}}$ $\Delta \textit{bla}_{\text{CTX-M-3}}$ $\Delta \textit{bla}_{\text{TEM-1}}$) strain. Colonies observed within the inhibition zone represent spontaneous subpopulations undergoing transient gene amplification (heteroresistance).

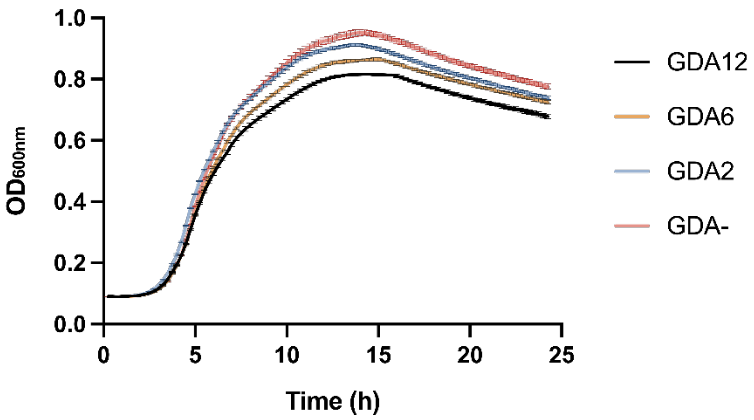


**Supplementary Fig. S3: Growth kinetics of GDA−, GDA2, GDA6, GDA12 strains in antibiotic-free medium.** The $OD_{600}$ values measured over 24 hours illustrate the overall growth performance and total biomass accumulation, demonstrating that strains with higher GDA copy numbers accumulate a lower final biomass compared to the GDA− strain. Data were presented as means ± S.D from three independent biological replicates.

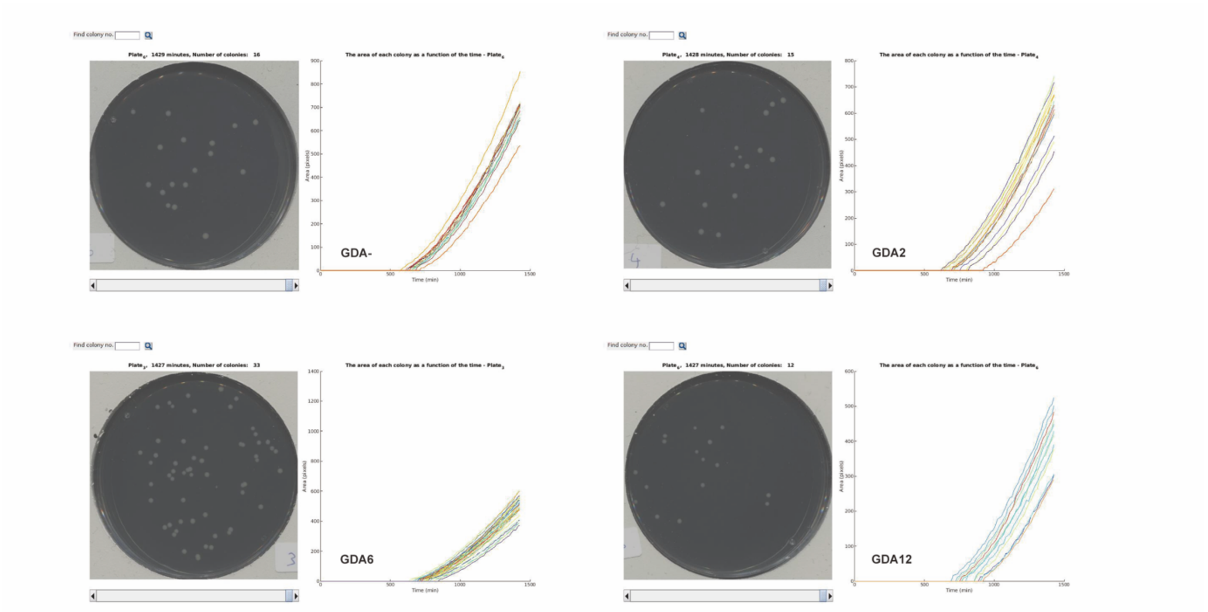


**Supplementary Fig. S4: Representative ScanLag analysis output for GDA−, GDA2, GDA6, and GDA12 strains under antibiotic-free conditions.** The plots display the growth dynamics of individual colonies on antibiotic-free agar plates. For each strain, the left panel displays the processed plate image identifying individual colonies, while the right panel plots the colony size (pixels) incubation time (minutes) of each detected colony. Each curve corresponds to a single colony. This analysis assesses the heterogeneity in lag time and growth rates among the population.

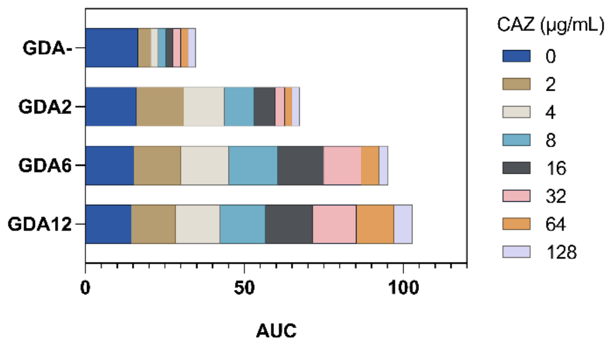


**Supplementary Fig. S5: Area Under the Curve (AUC) analysis of bacterial growth under increasing antibiotic concentrations.** The total growth of strains GDA−, GDA2, GDA6, and GDA12 was quantified as AUC over a period of 24 hours in the presence of ceftazidime ranging from 0 to 128 µg/mL. The plot illustrates the dose-dependent growth inhibition and distinct resistance profiles of the variants. Data are shown as means ± S.D from three independent biological replicates.

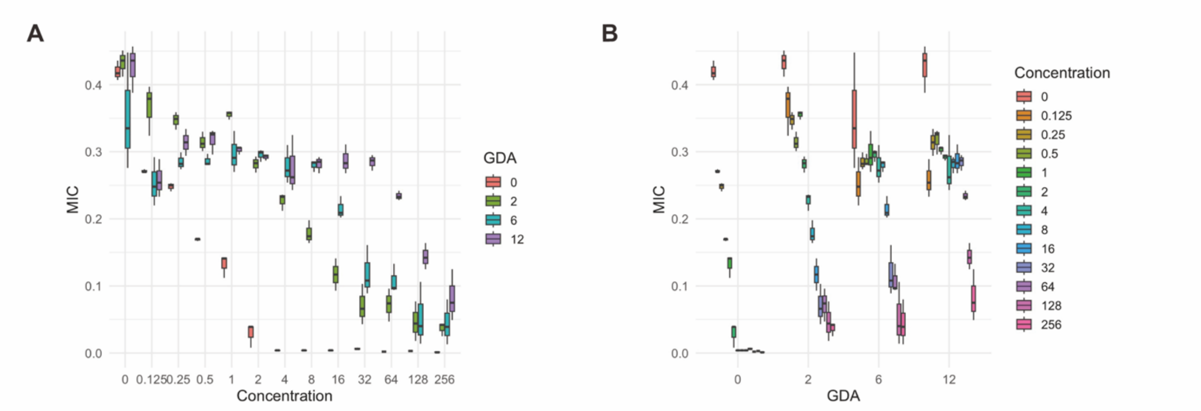


**Supplementary Fig. S6: Gene dosage shapes MIC distributions across antibiotic concentrations.** Boxplots summarize Minimum Inhibitory Concentration (MIC) values across GDA copy-number variants and ceftazidime concentrations (n = 3). **(A)** MIC distribution across antibiotic concentrations (x axis), grouped by GDA level (colors). **(B)** MIC distribution across GDA levels (x axis), grouped by antibiotic concentration (colors). Center lines indicate medians; box bounds indicate the interquartile range (IQR).

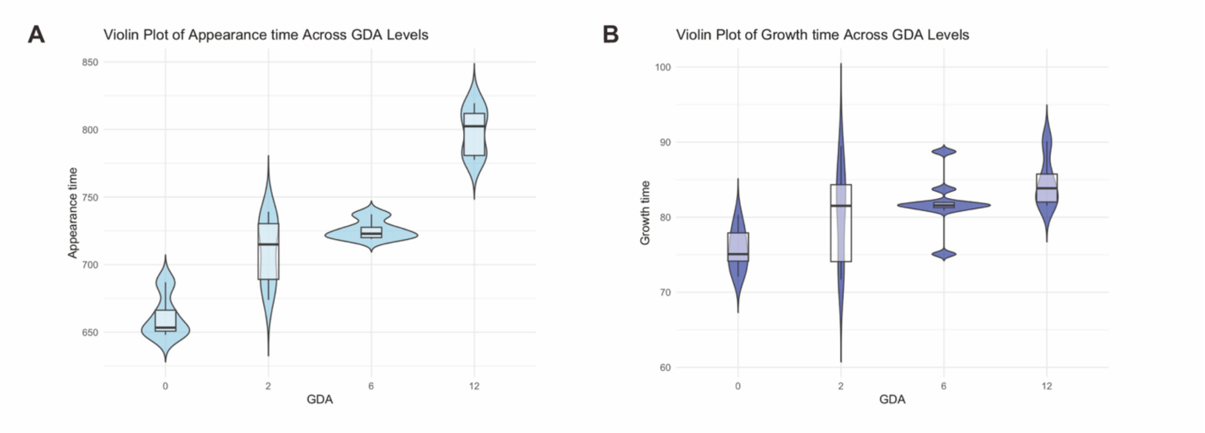


**Supplementary Fig. S7: Distributions of ScanLag-derived colony growth parameters across GDA copy-number variants.** Violin plots show the distribution of **(A)** colony appearance time and **(B)** colony growth time for each GDA-fixed strain (n = 3 independent biological replicates; total colonies analyzed = 9). The violin shapes represent kernel density estimates; embedded boxplots indicate the median and interquartile range (IQR).

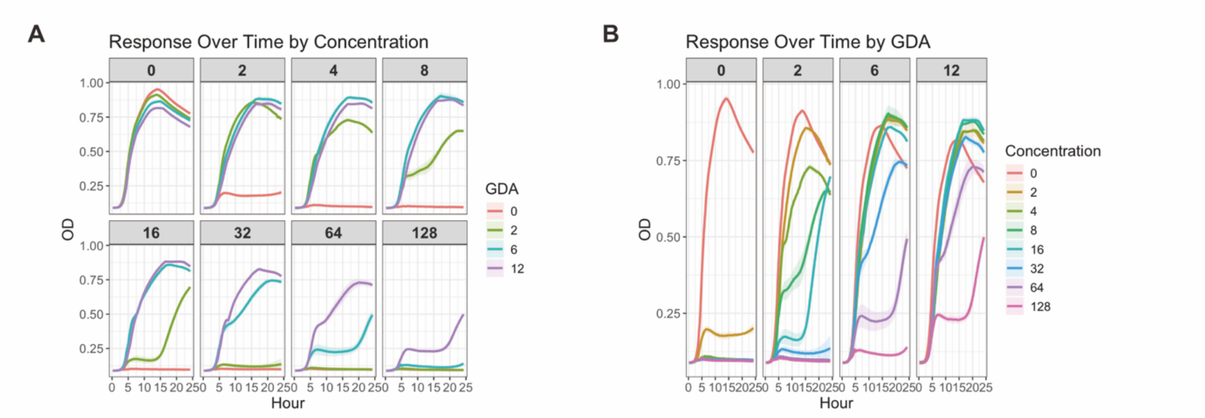


**Supplementary Fig. S8: Growth-curve dynamics across CAZ gradients and GDA copy-number variants.** Smoothed growth curves (OD600 over 24 h) are shown for GDA−, GDA2, GDA6, and GDA12 strains across ceftazidime (CAZ) concentrations ranging from 0 to 128 μg/mL (n = 3). **(A)** Growth curves faceted by CAZ concentration; colors denote GDA level. **(B)** Growth curves faceted by GDA level; colors denote CAZ concentration. Solid lines indicate mean fitted OD and shaded ribbons indicate 95% confidence intervals calculated from replicate variability (mean ± 1.96×SE).

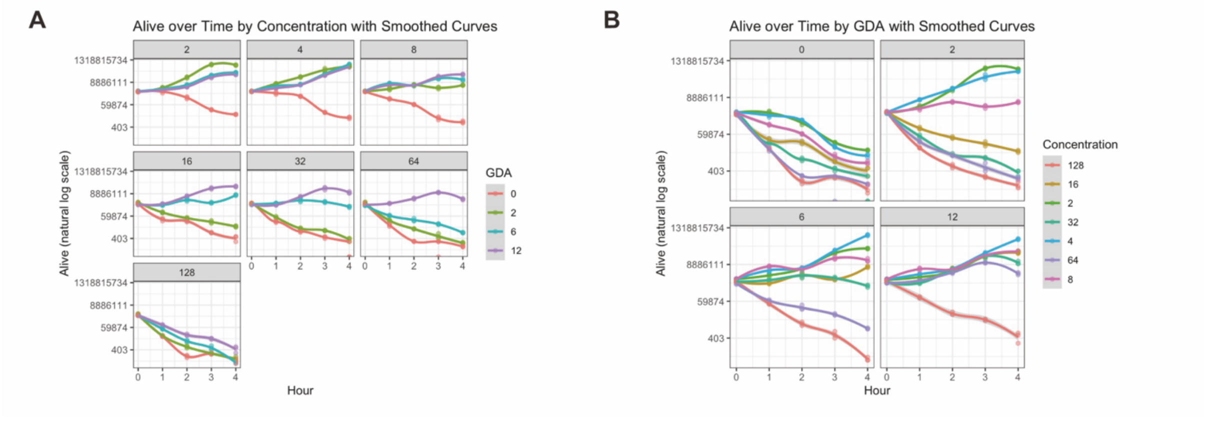


**Supplementary Fig. S9: Time-kill kinetics across CAZ gradients and GDA copy-number variants.** Time-kill assays for GDA−, GDA2, GDA6, and GDA12 were performed over 4 h across ceftazidime (CAZ) concentrations (n = 3). Points show viable counts (CFU/mL) measured at each time point and plotted on a log scale (y axis). Curves indicate LOESS-smoothed trends overlaid for visualization of killing dynamics. **(A)** Panels are faceted by CAZ concentration; colors denote GDA level. **(B)** Panels are faceted by GDA level; colors denote CAZ concentration.

**Supplementary Table S1: Model selection and comparison statistics for models fitted to Minimum Inhibitory Concentration (MIC) data.** Model Description denotes the model structure in R syntax notation (+ indicates main effects; * indicates interaction effects). Models with the lowest Akaike Information Criterion (AIC) values (indicating the best trade-off between goodness-of-fit and complexity) were selected for final analysis. df, degrees of freedom. GDA, gene duplication amplification copy number; Con, ceftazidime concentration; GDAF, GDA treated as a factor.

| Model | Description | df | AIC |
|---|---|---|---|
| mMic0 | Intercept only | 2 | -186.11 |
| mMic1 | Concentration * GDAF | 9 | -297.09 |
| mMic2 | Concentration + GDAF | 6 | -299.16 |
| mMic3 | Concentration | 3 | -242.61 |
| mMic4 | GDAF | 5 | -220.28 |

**Supplementary Table S2: Model selection and comparison statistics for models fitted to colony appearance time (ScanLag) data.** Model Description denotes the model structure in R syntax notation (~ indicates formula definition). Models with the lowest Akaike Information Criterion (AIC) values were selected for final analysis. df, degrees of freedom. GDA, gene duplication amplification copy number; GDA_num, GDA copy number treated as a continuous numeric variable.

| Model | Description | df | AIC |
|---|---|---|---|
| mAT0 | Intercept | 2 | 390.0798 |
| mAT1 | Time ~ GDA_num | 3 | 321.6055 |
| mAT1F | Time ~ GDA (factor) | 5 | 311.4911 |
| mAT2 | Time ~ GDA_num + $GDA\_num^2$ | 4 | 323.5527 |
| mAT0 | Intercept | 2 | 390.0798 |

**Supplementary Table S3: Model selection and comparison statistics for generalized additive models (GAMs) fitted to growth curve data.** Model Description denotes the model structure in R syntax notation (+ indicates main effects; s() indicates a smoothing function). Models with the lowest Akaike Information Criterion (AIC) values were selected for final analysis. df, degrees of freedom. GDA, gene duplication amplification copy number; Con, ceftazidime concentration; GDAF, GDA treated as a factor.

| Model | Description | df | AIC |
|---|---|---|---|
| GCGam0 | s(Hour) only | 9.68 | 1904.09 |
| GCGam1 | GDAF + ConF + s(Hour, by=interaction(GDAF,ConF)) | 162.17 | -14398.23 |
| GCGam2 | GDAF + ConF + s(Hour, by=GDAF) + s(Hour, by=ConF) | 65.56 | -11641.37 |
| GCGam3 | GDAF + ConF + s(Hour, by=GDAF) | 38.5 | -8493.71 |
| GCGam3.1 | GDAF + s(Hour, by=GDAF) | 28.79 | -2245.21 |
| GCGam3.2 | s(Hour, by=GDAF) | 23.97 | 1041.28 |
| GCGam4 | GDAF + ConF + s(Hour, by=ConF) | 58.09 | -8528.24 |
| GCGam4.1 | ConF + s(Hour, by=ConF) | 46.74 | -2869.56 |
| GCGam4.2 | s(Hour, by=ConF) | 31.6 | 1045.15 |
| GCGam0c | s(Hour) | 9.68 | 1904.09 |
| GCGam1c | s(Hour, GDA) | 28.27 | -2249 |
| GCGam2c | s(Hour, Con) | 30.19 | -2771.69 |
| GCGam3c | s(Hour, GDA, Con) | 110.69 | -27408.52 |

**Supplementary Table S4: Model selection and comparison statistics for models fitted to time-kill kinetics data.** Model Description denotes the model structure in R syntax notation (+ indicates main effects; * indicates interaction effects). Models with the lowest Akaike Information Criterion (AIC) values were selected for final analysis. df, degrees of freedom. GDA, gene duplication amplification copy number; Con, ceftazidime concentration.

| Model | Description | df | AIC |
|---|---|---|---|
| m0TKall | Intercept | 2 | 2390.42 |
| m1TKall | Hour * GDA * Con | 29 | 1698.57 |
| m2TKall | Hour + GDA * Con | 16 | 2026.43 |
| m3TKall | Hour * GDA + Con | 11 | 1956.4 |
| m4TKall | Hour * GDA + GDA * Con | 17 | 1919.51 |
| m5TKall | Hour + GDA + Con | 10 | 2052.65 |
| m6TKall | Hour * GDA | 5 | 2239.74 |
| m7TKall | Hour + GDA | 4 | 2289.24 |
| m8TKall | GDA * Con | 15 | 2058.37 |
| m9TKall | GDA + Con | 9 | 2081.74 |
| m10TKall | Hour | 3 | 2378.28 |
| m11TKall | GDA | 3 | 2304.73 |
| m12TKall | Con | 8 | 2222.93 |

**Supplementary Table S5: Primer used in this study.**

| Primer name | Sequence 5´- 3´ | Application |
|---|---|---|
| p49658blaDHAH1P1 | TGAATCTGACGATACTTGCCGCCGTTACTCAC ACACGGAAGGTTAATTCTGATGTGTAGGCTGG AGCTGCTTCGA | *bla*$_{\text{DHA-1}}$ gene deletion mutagenesis |
| p49658blaDHAH2P2 | TACGGCCCCGGCGTATCCGCAGGGGCCTGTT CAGGAAAAAAATTATTCCAGCATATGAATATCC TCCTTAG | *bla*$_{\text{DHA-1}}$ gene deletion mutagenesis |
| p49658_blaDHA_Test_F | GCATGGGTGACATTCAGCTCAAT | PCR |
| p49658_blaDHA_Test_R | AGCTGTCAGTGCCCGATACTC | PCR |
| blaACT_H1P1 | CTGACGGGCCCGGACATCCCCTTGACTCGCT ATTACGGAAGATTACTGATGTGTGTAGGCTGG AGCTGCTTCGA | *bla*$_{\text{ACT-16}}$ gene deletion mutagenesis |
| blaACT_H2P2 | CGTTGTGTAGGCCGGGTAAGCGCAGCGCCAC CCGGCAATGTTTTACTGTAGCATATGAATATCC TCCTTAG | *bla*$_{\text{ACT-16}}$ gene deletion mutagenesis |
| blaACT_testF | TAACCGTTTGTCAGGCACAG | PCR |
| blaACT_testR | AGATGACAGCAGGGAAAATGC | PCR |
| blaCTX_H1P1 | GACTATTCATGTTGTTGTTATTTCGTCTCTTCC AGAATAAGGAATCCCATGTGTGTAGGCTGGAG CTGCTTCGA | *bla*$_{\text{CTX-M-3}}$ gene deletion mutagenesis |
| blaCTX_H2P2 | ATAAACAAAAACGGAATGAGTTTCCCCATTCC GTTTCCGCTATTACAAACCCATATGAATATCCT CCTTAG | *bla*$_{\text{CTX-M-3}}$ gene deletion mutagenesis |
| blaCTX_testF | CATTCTGGCGACGTCCGTATTT | PCR |
| blaCTX_testR | TCGTGGTGCTGAATTTTGACG | PCR |
| blaTEM_H1P1 | AGACAATAACCCTGGTAAATGCTTCAATAATAT TGAAAAAGGAAGAGTATGTGTGTAGGCTGGAG CTGCTTCGA | *bla*$_{\text{TEM-1}}$ gene deletion mutagenesis |
| blaTEM_H2P2 | CACCCCCCAATTATTTCAGTATGAGTAAACTTG GTCTGACAGTTACCAATGCATATGAATATCCTC CTTAG | *bla*$_{\text{TEM-1}}$ gene deletion mutagenesis |
| blaTEM_testF | GTGGCACTTTTCGGGGAAATGT | PCR |
| blaTEM_testR | TAGTGCTGATATTCAGGCCAC | PCR |
| 49658_RECAH1P1 | AGTCCATGGTGAAGCGCAGTTGCTTCTCCCGG CATGACAGGAGTAATAATGTGTGTAGGCTGGA GCTGCTTCGA | *recA* gene deletion mutagenesis |
| 49658_RECAH2P2 | CAGCAGCCCTTCATTTTTATCCGAGAGGATTAA AAGTCTTCGTTGGTTTCCATATGAATATCCTCC TTAG | *recA* gene deletion mutagenesis |
| 49658_RECAF | CACTTGATACTGTATGACTATACA | PCR |
| 49658_recAR2 | ACGCACGGCAAGAATACG | PCR |
| blaDHA-1 forward | GTTCAGCCGTATAACCGTGCTTC | qPCR |
| blaDHA-2 reverse | TCACAATCGCCACCTGTTTTTCC | qPCR |
| trpA (EC) primer forward | CTACGACAACGCATTTGCACAAC | qPCR |
| trpA (EC) primer reverse | TCGATAATCTTCAGCGACTGCTCC | qPCR |
